# Absent expansion of pericentral hepatocytes and altered physiology in Axin2CreERT2 mice

**DOI:** 10.1101/2020.12.09.405936

**Authors:** Stephanie May, Miryam Müller, Callum R Livingstone, George Skalka, Colin Nixon, Ann Hedley, Robin Shaw, William Clark, Thomas M Drake, Christos Kiourtis, Ana Sofia Rocha, Owen J Sansom, Thomas G Bird

**Author notes:** Contact info Dr Thomas G. Bird, Cancer Research UK Beatson Institute, Glasgow, G61 1BD, UK.

## Abstract

Understanding how the liver regenerates is a key biological question. Hepatocytes are the principle regenerative population in the liver. Recently, numerous lineage tracing studies (which apply genetic tagging to a restricted population and track its descendants over time) have reported conflicting results using a variety of hepatocyte based reporting systems in mice^1,2^. The first significant lineage tracing from a distinct subpopulation of hepatocytes in homeostasis reported hyper-proliferation of self-renewing pericentral hepatocytes with their subsequent expansion across the liver lobule^3^. This study used a CreERT2 construct knocked into the endogenous *Axin2* locus; here termed Axin2CreERT2. Subsequent studies, using either a different pericentral marker (Lgr5^4^) or a different AxinCreERT2 transgene^5^, did not show lineage tracing. Here we aim to reconcile these discrepancies by re-evaluating lineage tracing in the Axin2CreERT2 knock-in model and explore the physiological consequences of this mutant allele. We were unable to find evidence of expansion of an Axin2CreERT2 labelled population and show that this population, whilst zonated, is spread throughout the lobule rather than being zonally restricted. Finally, we report that this allele results in profound perturbation of the Wnt pathway and physiology in the mouse.

## Results

### Absence of expansion from an Axin2^+^ population in homeostasis

In order to define a baseline labelled population in the Axin2CreERT2^+/WT^ mouse model we administered mice with 4mg tamoxifen, which does not affect Wnt target gene expression in WT mice (Supplementary Fig. 1a,b). To facilitate the registration of labelling to specific cells responsive to the Axin2CreERT2 we used an intracellular LSL-RFP reporter. We analysed labelling efficiency at 3 time points within a week after induction (Fig. 1a). We observed maximal labelling of pericentral hepatocytes after 5 days (Fig 1b,c), with significantly less labelling in male compared to female mice. For simplicity, we hereon refer to quantitative data for female mice, but trends were similar in both sexes. Next, we studied the zonal restriction of hepatocyte labelling in this model using hepatic zones defined by expression of glutamine synthetase (GS) and E-cadherin (Fig. 1d). Zonal analysis showed a progressive increase in labelling in zone 3 from day 2, plateauing from day 5 to day 7 when 35% of hepatocytes in zone 3 were labelled (Fig. 1e). However, only approximately half of all labelled hepatocytes were contained within zone 3 (Supplementary Fig. 1c); itself comprising consistently 6-8% of all hepatocytes (Supplementary Fig. 1d). In the remaining liver lobule, 1.5-3.5% of hepatocytes were also labelled (Fig. 1b open arrows, Supplementary Fig. 1e), including hepatocytes within zone 1 (Fig. 1f,g); albeit to a lesser extent than in other liver zones (Supplementary Fig. 1f). Consistent with recombination of the reporter in Axin2CreERT2^+/WT^ across the lobule, *Axin2* was expressed throughout the lobule in WT mice but with a highly graded expression from zones 3 to 1 (Fig. 1h). Our data suggests that, in this knock-in reporter system, CreERT2 also has graded expression across the lobule. Therefore, whilst this labelling system preferentially labels zone 3 hepatocytes, the labelling is not restricted to the pericentral hepatocytes, as previously reported^3^, and indeed the bulk of labelling occurs outside zone 3. Furthermore, as incomplete labelling occurs at early time points, 5-7 days (but not as early as 2 days) is a more appropriate baseline for lineage tracing in this model.

**Figure 1:**
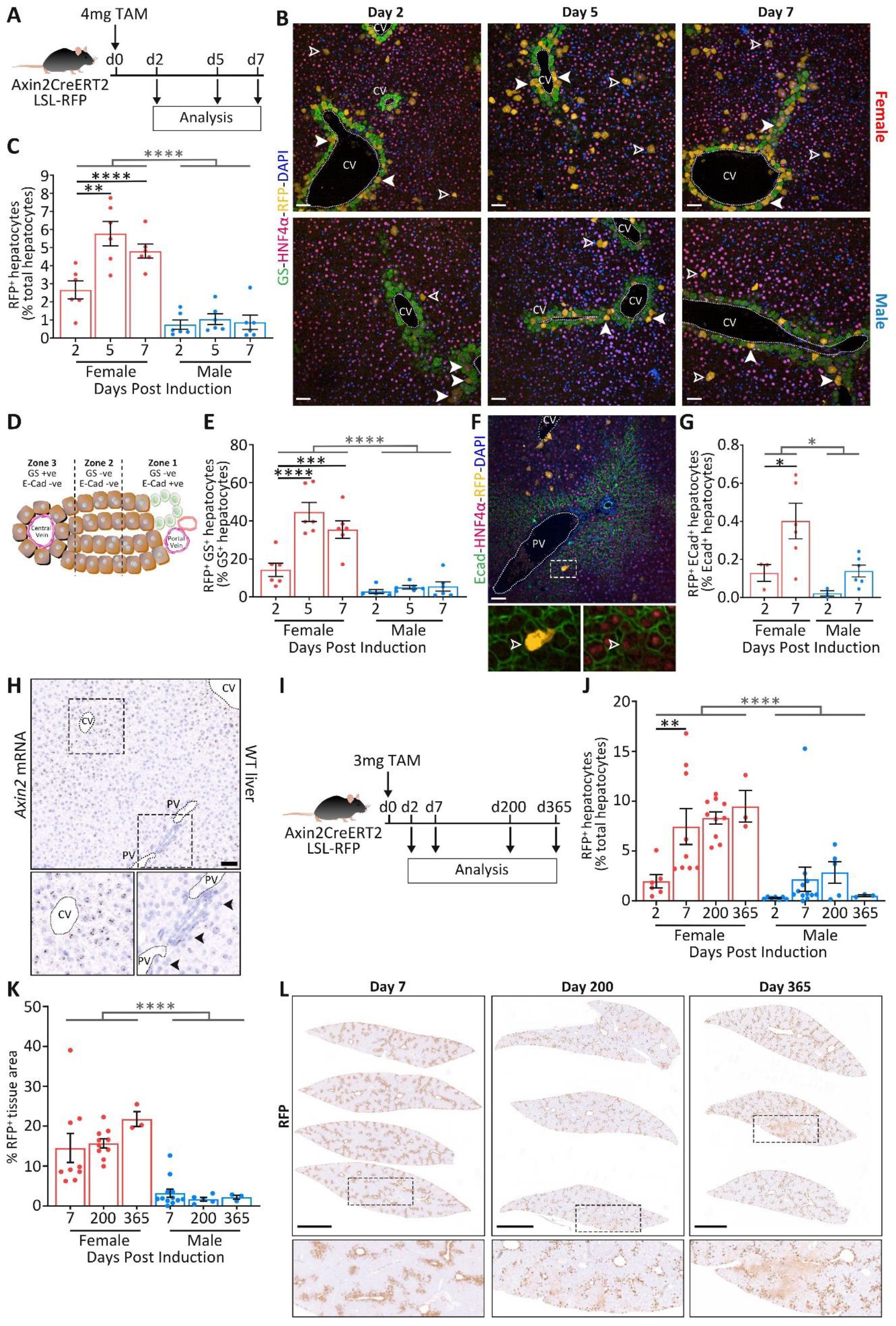
Axin2^+^ hepatocytes do not preferentially contribute to homeostatic liver regeneration. (**A**) Axin2CreERT2^+/WT^ progeny of Axin2CreERT2^+/WT^ mice crossed with LSL-RFP^+/+^ reporter mice were induced with 4mg Tamoxifen and analysed at 2, 5 and 7 days post induction. (**B**) Representative microscopic images highlighting central perivenular (GS; green), labelled (RFP; orange), hepatocytes (HNF4α; red) and nuclei (DAPI; blue) are shown for male and female mice at each time point post tamoxifen induction; demonstrating preferential labelling of hepatocytes around the central vein (CV, solid arrows) but also hepatocytes outside of this area (open arrows). Dashed white line = border of CV, scale bar = 50μm. (**C**) Quantification of RFP^+^ hepatocytes over the time course compared to day 2; 21-44 (median = 31) high power fields in N = 6 mice at each time point separated by sex, two-way ANOVA, mean ± SEM, total hepatocytes 219,768/225,318/228,551 quantified at days 2/5 and 7 respectively. (**D**) We defined hepatocytes in liver zones as follows; zone 1 (periportal region) GS^-^/Ecad^+^, zone 2 GS^-^/Ecad^-^ and zone 3 (pericentral region) GS^+^/Ecad^-^. (**E**) Quantification of the proportion of zone 3 hepatocytes that are RFP^+^, N = 6 mice at each time point, two-way ANOVA, mean ± SEM. (**F**) Microscopic images showing periportal (Ecad; green), labelled (RFP; orange), hepatocytes (HNF4α; red), in a female mouse 2 days post induction; dashed white line = border of CV and portal vein (PV), open arrow highlights a RFP labelled hepatocyte within the Ecad^+^ zone 1, scale bar = 50μm. (**G**) Quantification of the proportion of zone 1 hepatocytes labelled with RFP^+^ over time post induction; 11-20 (median = 15) high power fields in N = 3 and 6 mice at days 2 and 7 respectively for both sexes, two-way ANOVA, mean ± SEM, total hepatocytes 49,668/107,642 quantified at days 2 and 7 respectively. (**H**) RNA *in situ* hybridisation for *Axin2* in WT murine liver demonstrating a zonated pattern of *Axin2* expression across the hepatic lobule; arrows highlight periportal hepatocytes expressing *Axin2*, scale bar = 50μm. (**I**) Long-term lineage tracing for up to 1 year was performed post induction of Axin2CreERT2^+/WT^ LSL-RFP^+/-^ mice with 3mg tamoxifen. (**J**) Quantification of RFP^+^ hepatocytes over time compared to day 7 baseline; 17-69 (median = 30) high power fields analysed in N ≥ 3 mice at each time point separated by sex, total hepatocytes 277,826/315,421/209,895/81,280 quantified at days 2/7/200 and 365 respectively, see Supplemental Fig.2 for examples of cell registration. No expansion from day 7 to either 200 or 365 days post induction; p values = 0.8951/0.6480 (day 200) and 0.8124/0.9699 (day 365) in males/females respectively; two-way ANOVA, mean ± SEM. N numbers at days 2, 7, 200 and day 365 were 7/6, 12/9, 5/10 and 3/3 for males/females respectively. (**K**) Additional quantification of RFP^+^ expansion from 7 days as quantified by tissue area in whole liver tissue areas; numbers at days 7, 200 and 365 were 12/9, 5/10 and 3/3 mice for males/females respectively. P values versus day 7 = 0.8561/0.8739 (day 200) and 0.9536/0.1201 (day 365) in males/females respectively; two-way ANOVA, mean ± SEM. (**L**) Representative low power microscopic images of whole lobes from individual mice stained with RFP are shown for day 7, 200 and 365; scale bars = 2.5mm. P = *<0.05; **<0.01; ***<0.001; **** <0.0001. Statistical comparisons between female vs. male are reported as the interaction factors from two-way ANOVA.

To further confirm labelling outside zone 3 we performed a depletion experiment by destroying the GS zone using the hepatotoxin carbon tetrachloride (CCl_4_). Here, we increased the labelling of hepatocytes using a repeated tamoxifen dosing regimen with CCl_4_ administered one week after the start of induction (Supplementary Fig. 1g). 24hrs post CCl_4_ there was efficient destruction of zone 3 hepatocytes but only a partial depletion of RFP labelled hepatocytes (Supplementary Fig. 1h,i). Labelled hepatocytes were preserved into the recovery phase with some reconstituted GS hepatocytes bearing the reporter 4 days post CCl_4_ (Supplementary Fig. 1j). Therefore, using a functionally defined zone, we again find evidence of genetic labelling outside zone 3.

For long-term lineage tracing studies, with the rationale of maximising restriction of hepatocyte labelling to zone 3, we used lower doses of tamoxifen (Fig. 1i). The 3mg tamoxifen labelling dose achieved comparable labelling of hepatocytes (~7%; Fig. 1j) to that previously published in this model at day 7 using 4mg^3^. Again we observed progressive labelling from day 2 to 7 and therefore used day 7 as a baseline. At day 7 no areas of confluent hepatocyte labelling were observed. Lineage tracing was then performed in otherwise untreated mice for up to 1 year. During this time no expansion of the labelled hepatocyte population occurred in either female or male mice when evaluating either hepatocytes or total labelled liver area (Fig. 1j-l, Supplementary Fig. 2). Some patches of confluent labelling of hepatocytes were observed at 200 and 365 days. These were infrequent (approximately 3/ lobe) and occurred overwhelmingly in female mice. Therefore, whilst some localised expansion events are observed, there was no overall expansion of the labelled population in Axin2CreERT2^+/WT^ mice.

**Figure 2:**
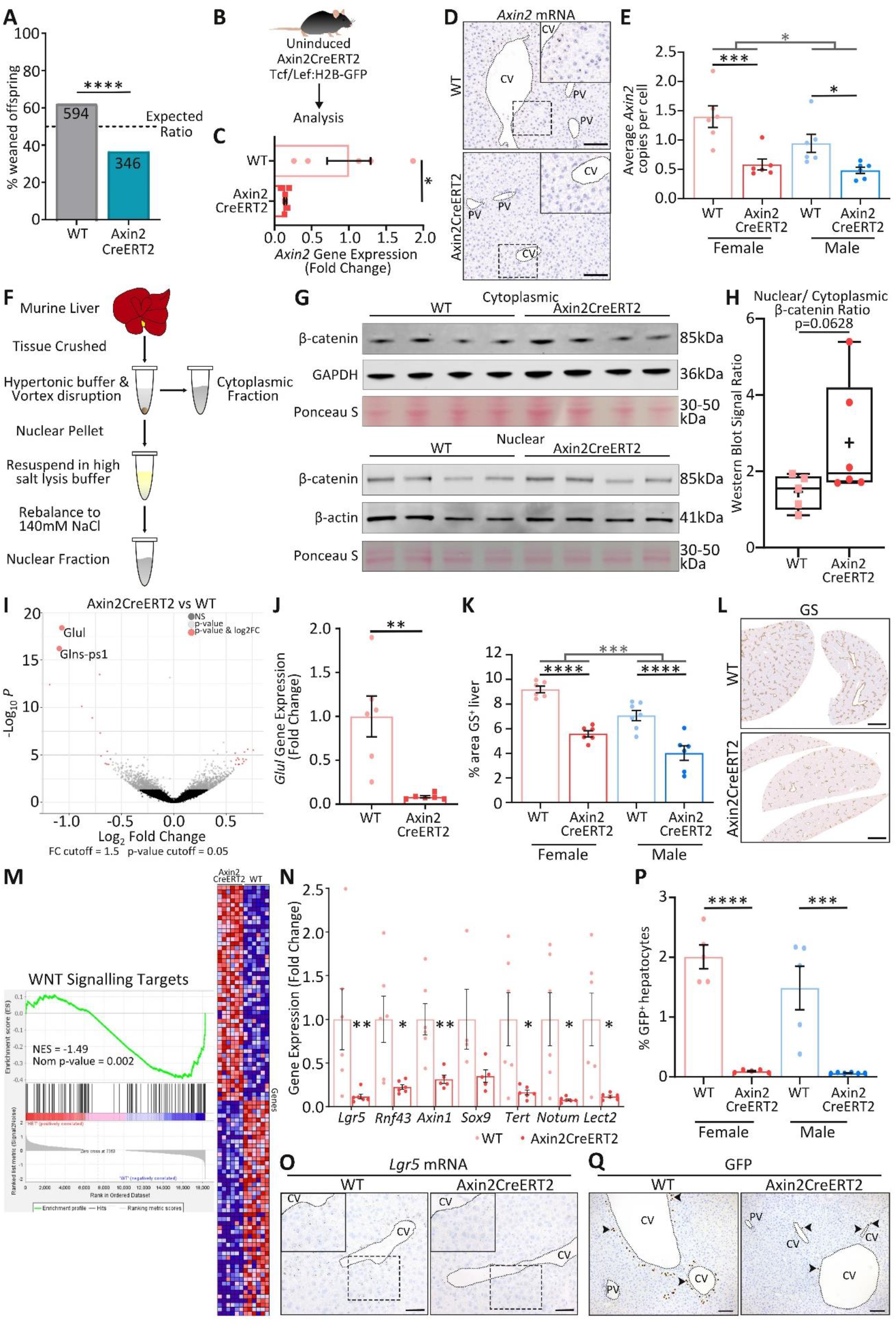
Axin2CreERT2 knock-in construct alters liver physiology. (**A**) Expected Mendelian ratios of offspring weaned were not observed from WT x Axin2CreERT2^+/WT^ mating’s (940 mice from 57 mating’s over 4 years); chi-squared test. (**B**) For all homeostatic experiments, tissue was harvested from uninduced Axin2CreERT2 wild type (referred to as WT) and heterozygous mice, all with a Tcf/lef:H2B-GFP^+/-^ reporter. (**C**) *Axin2* gene expression from whole liver by qRT-PCR in female mice; N = 5 WT mice and 6 Axin2CreERT2^+/WT^ mice, t-test, mean ± SEM. (**D**) Microscopy of RNA *in situ* hybridisation for *Axin2* in female livers comparing uninduced Axin2CreERT2^+/WT^ versus WT, with quantification in (**E**); scale bar = 100μm, N = 6 per cohort, two-way ANOVA, mean ± SEM. (**F**) Schematic for isolating nuclear and cytoplasmic protein fractions. (**G**) Representative western blots of β-catenin in cytoplasmic and nuclear fractions from uninduced WT and Axin2CreERT2^+/WT^ livers. GAPDH and β-actin are the loading controls for the cytoplasmic and nuclear protein fractions respectively. (**H**) Quantification of nuclear to cytoplasmic β-catenin ratio in uninduced WT and Axin2CreERT2^+/WT^ mice, each fraction is normalised to β-actin and GAPDH respectively; N = 5 WT mice and 6 Axin2CreERT2^+/WT^ mice, one-tailed Mann-Whitney test, box plot with centre line showing the median values (interquartile range (IQR)), for WT 1.55 (1.00-1.88) and Axin2CreERT2^WT/-^ 1.95 (1.71-4.20), whiskers represent minimum and maximum values, ‘+’ = mean. (**I**) Volcano plot displaying differentially expressed genes from RNAseq analysis of whole female livers of uninduced Axin2CreERT2^+/WT^ compared to WT mice; N = 6 per cohort. (**J**) *Glul* gene expression validated from whole liver by qRT-PCR in female mice, N = 6, two-tailed Mann-Whitney test, mean ± SEM. (**K**) Quantification of GS positivity by tissue area in whole liver from male and female mice, with representative low power microscopic images of whole lobes from individual female mice (**L**); N = 7/6 and 6/6 in WT/Axin2CreERT2^+/WT^ mice for males and females respectively, two-way ANOVA, mean ± SEM, scale bars = 1.5mm. (**M**) Gene set enrichment analysis (GSEA) plot demonstrating down regulation of Wnt signalling target genes in uninduced female Axin2CreERT2^+/WT^ mice compared to WT and respective heat map of the top 50 features for each genotype, N = 6 mice per cohort. Hits mark the position of genes in published datasets. (**N**) Gene expression of Wnt target genes from female whole livers of uninduced WT and Axin2CreERT2^+/WT^ mice by qRT-PCR; N = 4-6 mice, two-tailed Mann-Whitney (or t-tests if normally distributed), mean ± SEM. (**O**) Microscopy of RNA *in situ* hybridisation for *Lgr5* in a female liver demonstrating a reduction in uninduced Axin2CreERT2^+/WT^ mice compared to WT mice; scale bars = 100μm. (**P**) Using the Tcf/lef:H2B-GFP as a canonical Wnt pathway reporter GFP^+^ hepatocytes were quantified from IHC sections stained with GFP in uninduced WT and Axin2CreERT2^+/WT^ mice, with representative microscopic images (**Q**); N = 5 mice per cohort, except for male Axin2CreERT2^+/WT^ mice where N = 6, two-way ANOVA, mean ± SEM, scale bars = 100μm. P = *<0.05; **<0.01; ***<0.001; **** <0.0001. Statistical comparisons between female vs. male are reported as the interaction factors from two-way ANOVA.

Given the previous report of increased proliferation in zone 3, we compared proliferation in WT mice using both young and aged mice of both sexes. Consistent with other reports^4^, we did not find hyper-proliferation of cells within zone 3 (Supplementary Fig. 1k).

### Wnt pathway is altered in homeostasis in the presence of the Axin2CreERT2 allele

As Axin2 is part of the Wnt/β-catenin pathway we hypothesised that this knock-in allele of Axin2CreERT2 may cause a haploinsufficiency phenotype. We began by examining the physiological effects of the presence of this allele in uninduced mice. Breeding Axin2CreERT2^+/WT^ mice, we were unable to generate homozygote pups (0/14 at weaning; with 4 dying perinatally). In heterozygote x WT mating’s we obtained significantly fewer heterozygote pups at weaning than predicted (Fig. 2a, Supplementary Fig. 3a). Despite this, uninduced heterozygote mice assessed during adulthood have equivalent body weight (data not shown), relative liver weight (Supplementary Fig. 3b), hepatocyte proliferation (Supplementary Fig. 3c) and liver biochemistry (Supplementary Fig. 3d-f) to WT littermates.

Next, we assessed the Wnt pathway in Axin2CreERT2^+/WT^ mice. Here we used Wnt reporter mice (Tcf/Lef:H2B-GFP, Fig. 2b) expressing GFP in Wnt/β-catenin responsive cells^6^. *Axin2* expression was reduced in uninduced Axin2CreERT2^+/WT^ mice (Fig. 2c-e, Supplementary Fig. 3g). Expression of *Axin2* was again zonated but not zonally restricted in both genotypes (Fig. 2d).

Given Axin2’s role in the β-catenin destruction complex we next assessed the Wnt pathway in the liver. Using cell fractionation (Fig. 2f) we observed a trend of increased nuclear β-catenin in Axin2CreERT2^+/WT^ mice compared to WT littermates (Fig. 2g,h, Supplementary Fig. 3h,i) suggestive of hyper-activation of the Wnt/β-catenin pathway. We then examined Wnt/β-catenin pathway activation. First, in an unbiased approach, we performed transcriptomic analysis. We found numerous dysregulated genes (Fig. 2i, Supplemental Fig. 3j), with expression of Wnt pathway targets GS (*Glul*) and a pseudogene of GS (*Glns-ps1*) being the most altered in uninduced Axin2CreERT2^+/WT^ mice compared to WT mice. A reduction of GS was confirmed by qRT-PCR (Fig. 2j) and at the protein level (Fig. 2k,l). Additionally, a reduction was seen across a larger panel of Wnt pathway transcriptional targets (Fig. 2m-o, Supplementary Fig. 3k). We sought to validate these findings with the *in vivo* Wnt pathway reporter system and found reduced Tcf/Lef driven GFP reporter in uninduced Axin2CreERT2^+/WT^ mice compared to WT littermates (Fig. 2p,q). Taken together, our data demonstrate altered physiology in mice possessing the Axin2CreERT2 allele with profound changes in the Wnt signalling pathway in the liver, particularly in female mice.

### Presence of the Axin2CreERT2 allele does not affect liver regeneration

In view of the important role of Wnt/β-catenin in liver regeneration^7^ and the dysregulation of this pathway in mice possessing the Axin2CreERT2 allele we next examined whether the regenerative response is altered in these animals. To address this we performed the archetypical 70% partial hepatectomy regenerative model in uninduced mice (Fig. 3a) followed by analysis at either peak regeneration (day 2) or at a recovery time point (day 7). This model causes global parenchymal loss rather than zonal specific damage caused by most hepatotoxins. Using the Tcf/Lef:H2B-GFP reporter we also investigated Wnt responses. Here, recovery of liver weight, indicative of regeneration post partial hepatectomy, was independent of genotype in both sexes (Fig. 3b,c). Peak and total hepatocyte proliferation responses were also equivalent (Fig. 3d,e). Tcf/Lef based reporting after partial hepatectomy was not altered in either control or Axin2CreERT2^+/WT^ mice (Fig. 3f). Finally, taking an unbiased approach we performed global hepatic transcriptomic analysis and found that highly similar transcriptional profiles occurred in both genotypes induced by partial hepatectomy (Fig. 3g-i) with no evidence of specific dysregulation of the Wnt/β-catenin pathway. Therefore, we conclude that, whilst differences in physiology and the Wnt/β-catenin pathway exist in homeostasis, we find no evidence of altered regenerative response in Axin2CreERT2^+/WT^ adult animals in response to partial hepatectomy.

**Figure 3:**
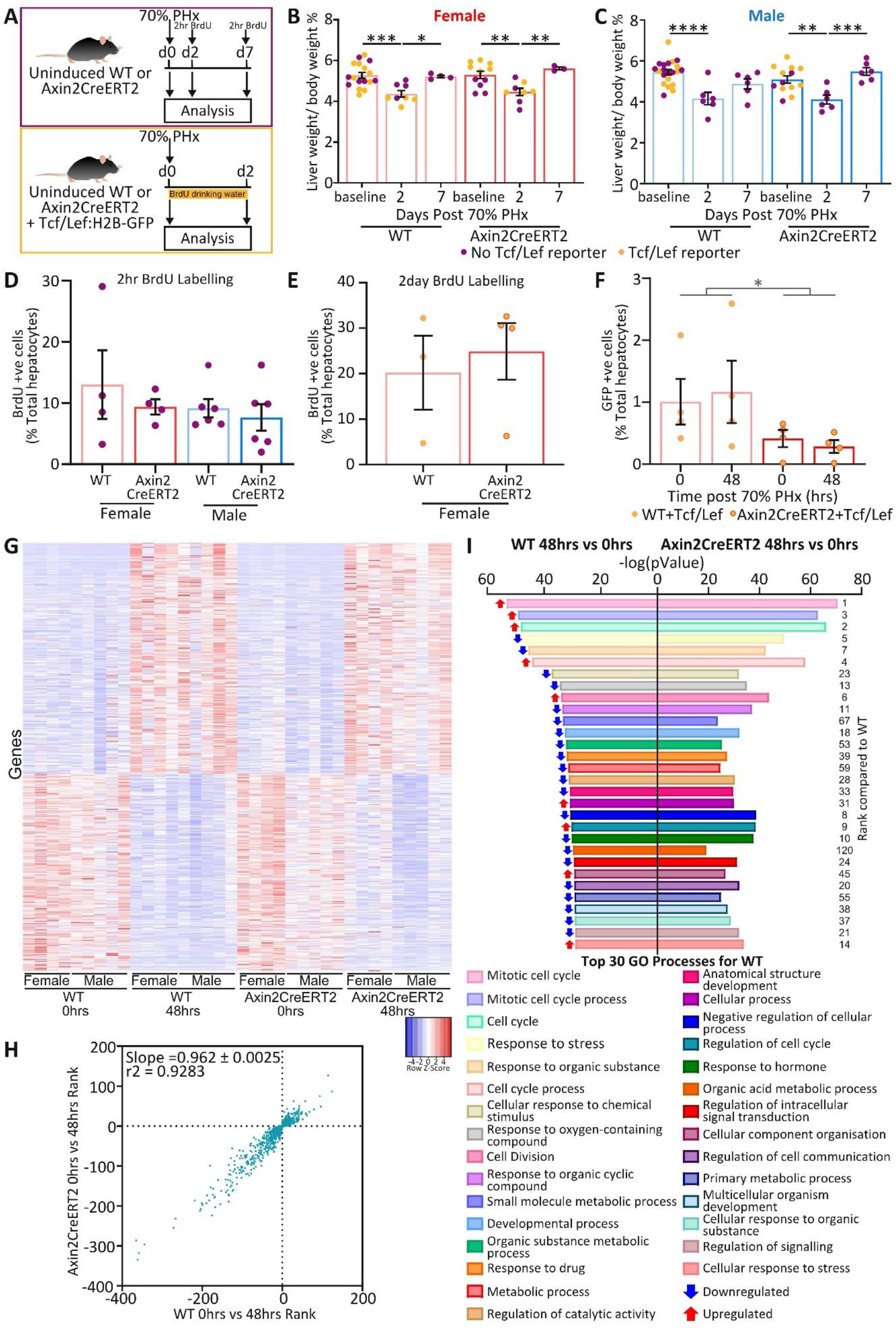
Liver regeneration is not affected by the presence of the Axin2CreERT2 construct. (**A**) Liver regeneration in response to 70% partial hepatectomy (PHx) in uninduced WT and Axin2CreERT2^+/WT^ mice (purple) or in mice also harbouring the Tcf/Lef:H2B-GFP reporter (orange). Mice were either administered BrdU 2 hours before sampling, or for 2 days in the drinking water after surgery respectively. Tissue was analysed at 2 or 7 days post-surgery. Liver weight to body weight ratios were calculated for (**B**) female and (**C**) male WT and Axin2CreERT2^+/WT^ mice at baseline and days 2 and 7 post PHx; N = 21/17, 6/8 and 6/4 in WT mice and 13/11, 6/8, and 6/3 in uninduced Axin2CreERT2^+/WT^ mice at baseline, day 2 and day 7 for males/females respectively, two-way ANOVA, mean ± SEM. Hepatocyte proliferation was quantified from IHC sections stained with BrdU at day 2 post-surgery following either 2hr BrdU pulse (**D**) or continuous application after surgery (**E**); N = 4 female mice and 6 male mice per cohort (D) and 3 WT and 4 Axin2CreERT2+/WT mice (E), two-way ANOVA and two-tailed Mann-Whitney respectively, mean ± SEM. (**F**) Percentage of GFP^+^ hepatocytes responding to the Tcf/lef:H2B-GFP Wnt reporter before and 2 days after 70% PHx were quantified in uninduced WT and Axin2CreERT2^+/WT^ mice from IHC sections stained with GFP; N = 4 mice per cohort, two-way ANOVA, mean ± SEM. (**G**) Heat map displaying differentially expressed genes identified by RNAseq analysis of whole livers from uninduced Axin2CreERT2^+/WT^ and WT mice that occur following 70% PHx. This is using paired resected (0hrs, during surgery) and regenerated tissues (48hrs post-surgery) from both sexes, N = 9 mice per cohort (4 female and 5 male). (**H**) Ranking of differentially expressed genes in WT and Axin2CreERT2^+/WT^ mice (combined sexes, N = 9 for each genotype (4 female and 5 male) based on their π-score; linear regression slope = 0.962 ± 0.0025). (**I**) Top 30 significant gene ontology (GO) cellular processes, ranked by p value, altered in WT tissue after 70% PHx. The equivalent GO processes are shown for uninduced Axin2CreERT2^+/WT^ mice and their respective ranking compared to WT; (N = 9 per cohort). Arrows show directionality of change following PHx (red = up, blue = down). P = *<0.05; **<0.01; ***<0.001; **** <0.0001. Statistical comparisons between female vs. male are reported as the interaction factors from two-way ANOVA.

## Discussion

This study helps to reconcile the conflicting hepatocyte lineage tracing studies. Consistent with previous reports^3^ we find evidence of expansion of some cells within the population labelled by Axin2CreERT2. However, there is no increased expansion of the population overall, particularly when taking into account the delayed labelling which occurs within the first week post tamoxifen. Hence, these rare expansion events are balanced by loss of other cells within the population. Our data argue against a dominant expansion of cells from within the pericentral zone in homeostasis as previously proposed. Our data are consistent with other lineage tracing studies using zonally enriched Wnt/β-catenin pathway members^4,5^. Despite theoretical differences in labelling due to the site of knock-in and loss of endogenous regulatory elements, we do not find discrepancies in lineage tracing between our Axin2CreERT2 and the published Axin2 transgene^5^. Whilst our study is unable to resolve the identity of those cells within the labelled population which do expand, it is consistent with previous reports of scattered cells with distinct and possibly fluctuant phenotypes capable of preferential expansion^8–12^. Due to the lack of specific labelling within a tightly geographically defined population we are unable to define whether the expansion events we observe occur from a pericentral origin. The pattern of the labelling however does imply that there are vascular relationships to the expansion as margins of confluent labelled areas typically associate with vascular territories.

This study furthermore highlights the impact of haploinsufficiency when using a genetic knock-in within a functional locus. It was previously acknowledged that such haploinsufficiency may have an impact upon phenotype but upon examining hepatocyte proliferation, no phenotypic alteration was demonstrated^3^. This is in agreement with our data both in homeostatic and regenerative proliferation. None the less by performing additional analyses we show profound changes, both in Axin2 and in the Wnt pathway which Axin2 regulates^13^, in the chronic *Axin2* haploinsufficient state in the Axin2CreERT2 heterozygotes that survive to adulthood. Notably in these animals we observed an uncoupling of canonical Wnt signalling. Reduced Axin2 is expected to lead to impaired β-catenin degradation and consequently high nuclear β-catenin resulting in overexpression of canonical Wnt pathway targets. In chronic *Axin2* haploinsufficiency the latter becomes uncoupled, with instead reduced expression of canonical Wnt pathway targets. Notably this effect is more prominent in female mice. Therefore it appears that there is significant compensation in the regulatory networks of Wnt signalling in those animals surviving to adulthood. We propose that this compensation underlies the normal proliferation in these animals in the chronic adapted state compared to the well documented dysfunctional regeneration observed after acute Wnt/β-catenin signalling perturbation^14^.

We find that the sex effects are also relevant to lineage tracing. We show that comparison between male and female mice is inaccurate with differences both in stable labelling of a baseline population and in lineage tracing being strongly influenced by sex. This may be related to greater baseline labelling and/ or heightened dysregulation of the Wnt pathway in females. Therefore, it is imperative to report sex of animals used in such studies, consistent with guidelines for reporting in animal studies^15^, and to compare within but not between the sexes.

## Supporting information

Supplemental Results and Methods

## Author contributions

SM performed the animal studies, partial hepatectomy, designed and performed experiments, analysed the data, wrote the manuscript and produced figures. MM designed experiments, performed animal experiments, administered anaesthetic during partial hepatectomy and provided discussion. CRL performed experiments and analysed data. GS performed and analysed the western blots. AH and RS performed bioinformatic analysis. WC performed RNA sequencing. CN performed IHC and ISH. TD performed partial hepatectomy. ASR and CK assisted with animal studies. OJS provided resources, discussion and acquired funding. TGB designed experiments, analysed data, wrote the manuscript and acquired funding. All authors read the manuscript and provided critical comments.

## Acknowledgements

SM, CK, AH, RS, WC and CN were funded by Cancer Research UK (Grant number: A17196). CRL was funded by a Medical Research Scotland Vacation Scholarship (Grant number: Vac-1195-2018). GS was funded by Cancer Research UK (Grant number: A29252). MM, ASR and TGB were funded by the Wellcome Trust (Grant number: WT107492Z) and CRUK HUNTER Accelerator Award (Grant number: A26813). TMD was funded by the Graham Paterson Bequest Endowment (Grant number: 141725-01). OJS is funded by Cancer Research UK (Grant number: A21139).

The authors would like to thank CRUK Beatson Institute’s histological services, biological services, molecular technology and bioinformatics services, central services, Beatson Advanced Imaging Resource (BAIR) (core funded by CRUK – A17196) and the Clinical Pathology Lab (University of Glasgow) for their assistance. We would like to thank J. P. Iredale for constructive discussion and Catherine Winchester for critical review of the manuscript.

## Competing interest statement

The authors declare no conflict of interest.

## Additional Information

Supplementary Information is available for this paper.

Correspondence and requests for materials should be addressed to Dr Tom G Bird.

## Data availability statement

The datasets generated during and/or analysed during the current study are available from the corresponding author on reasonable request.

## Code availability

Custom code or mathematical algorithms were not used in this manuscript.

## Notes

### Competing Interest Statement

The authors have declared no competing interest.

