## Supplemental Results and Methods for "Absent expansion of pericentral hepatocytes and altered physiology in Axin2CreERT2 mice"

#### Supplementary Information

##### Supplemental Figures

Sup Fig 1

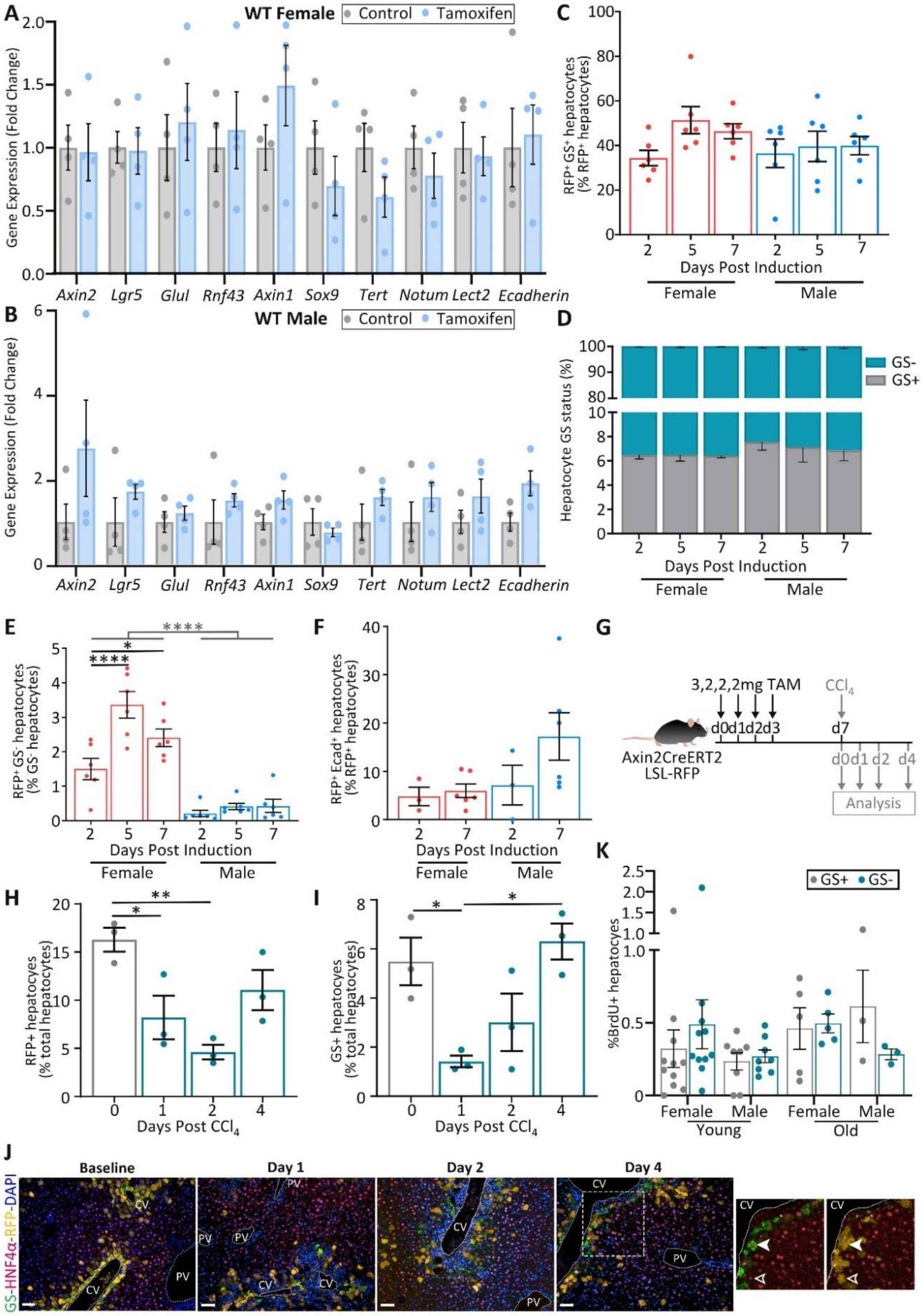

#### Sup Fig 1: Axin2CreERT2 lineage tracing in homeostasis and injury

qRT-PCR gene expression of Wnt target genes in liver from female (**A**) and male (**B**) C57Bl6 WT mice 2 days after i.p. administration of 4mg tamoxifen and uninduced WT controls sampled at the same time; N = 3-4 mice per cohort, two-tailed Mann-Whitney (or t-tests if normally distributed, all analyses were not significant), mean ± SEM. (**C**) Axin2CreERT2$^{+/WT}$; LSL-RFP$^{+/-}$ mice of both sexes were induced with 4mg tamoxifen and followed for 2, 5 and 7 days, see Fig. 1a. Quantification of the proportion of RFP$^{+}$ cells in Axin2CreERT2$^{+/WT}$ mice that were located within the pericentral region over the time course compared to day 2; N = 6 mice per cohort, two-way ANOVA, mean ± SEM. (**D**) Quantification of the proportion of the hepatic lobule that makes up the pericentral region (GS$^{+}$ zone 3) over the time course; N = 6 mice, two-way ANOVA, mean ± SEM. (**E**) Quantification of the proportion of hepatocytes outside zone 3 that are RFP$^{+}$ compared to day 2 post induction; N = 6 mice at each time point, two-way ANOVA, mean ± SEM. (**F**) Quantification of the proportion of labelled hepatocytes that are in zone 1 from IF stained sections for RFP, HNF4α and E-cadherin; N = 3 mice at day 2 and 6 mice at day 7, two-way ANOVA, mean ± SEM. (**G**) Female Axin2CreERT2$^{+/WT}$; LSL-RFP$^{+/-}$ mice were induced for four consecutive days with tamoxifen injections (3mg followed by 3 days of 2mg) and traced for 7 days from the first tamoxifen injection before subsequent induction of acute liver injury using carbon tetrachloride ($CCl_4$). Mice were harvested at baseline (day 0, no $CCl_4$) and days 1, 2 and 4 post $CCl_4$ administration. (**H**) Quantification of RFP$^{+}$ hepatocytes over the $CCl_4$ time course compared to baseline; 23-50 (median = 36) high power fields in N = 3 mice at each time point, one-way ANOVA, mean ± SEM, total hepatocytes 6,315/4,078/3,322/8,055 quantified at days 0/1/2 and 4 post $CCl_4$ respectively. (**I**) Administration of $CCl_4$ results in pericentral injury as quantified by GS$^{+}$ hepatocytes from IF stained liver sections; N = 3 mice at each time point, one-way ANOVA, mean ± SEM. (**J**) Microscopic images showing pericentral (GS; green), labelled (RFP; orange), hepatocytes (HNF4α; red) and nuclei (DAPI; blue), in a female mouse at baseline and 1, 2 and 4 days post $CCl_4$ administration; dashed white line = border of CV and PV, solid arrow indicates a GS$^{+}$/RFP$^{+}$ hepatocyte, open arrow indicates a GS$^{-}$/RFP$^{+}$ hepatocyte 4 days after $CCl_4$; scale bar = 50µm. (**K**) Levels of proliferation in homeostatic livers were quantified in young (8-12 weeks old) and aged (>1 year old) female and male WT mice from IHC sections stained with BrdU and split into GS$^{+}$ and GS$^{-}$ regions; N = 8/3 and 11/5 for young/old male and female mice respectively, two-way ANOVA, mean ± SEM. P = *<0.05; **<0.01; ***<0.001; **** <0.0001. Statistical comparisons between female vs. male are reported as the interaction factors from two-way ANOVA.

Sup Fig 2

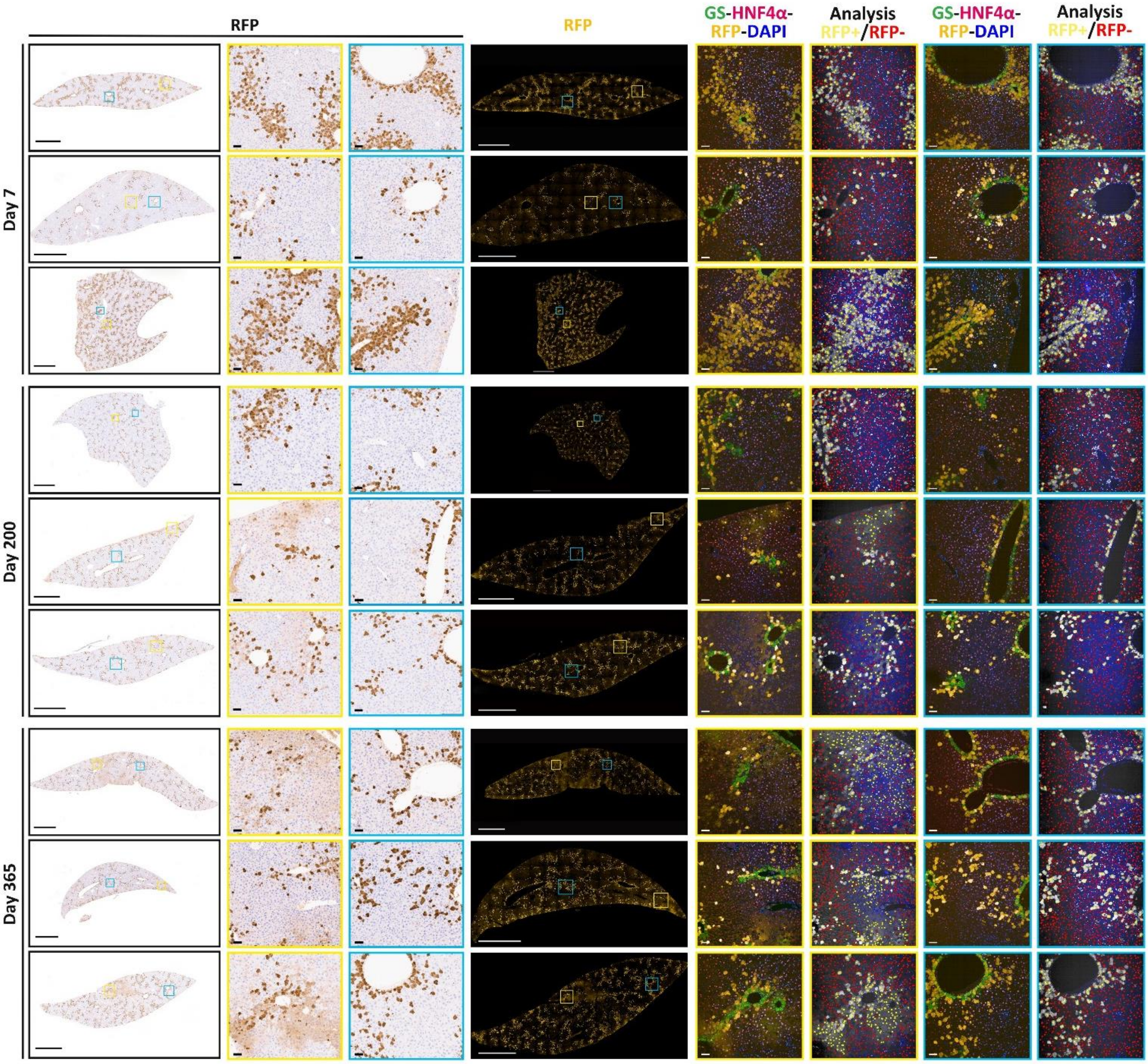


#### Sup Fig 2: Lineage tracing in Axin2CreERT2$^{+/WT}$; LSL-RFP$^{+/-}$ mice

Representative whole lobe images from N = 3 female Axin2CreERT2$^{+/WT}$; LSL-RFP$^{+/-}$ mice analysed for RFP labelling at 7, 200 and 365 days post induction with i.p. injection of 3mg tamoxifen. Serial sections with consecutive IHC and IF staining were performed for RFP (IHC) and GS, HNF4α, RFP and DAPI (IF) for quantification of RFP positivity using HALO or Columbus respectively. Analysis fields highlight hepatocyte recognition and which hepatocytes were quantified and registered as RFP$^{+}$ (yellow) or RFP$^{-}$ (red) using the Columbus cellular imaging and analysis platform. Rare diffuse confluent RFP labelling can be seen in some livers at 200 and 365 days post induction, but are accurately detected by RFP registering. Scale bars = 2mm for whole lobe and 50µm for field of view. Note: some of these images are also used in Fig. 1. Fig. 1L day 7 2nd lobe down = Sup Fig. 2 day 7 1st image flipped vertically. Fig. 1L day 200 3rd lobe down = Sup Fig. 2 day 200 3rd image flipped horizontally. Fig. 1L day 365 2nd lobe down = Sup Fig. 2 day 365 3rd image flipped horizontally.

Sup Fig 3

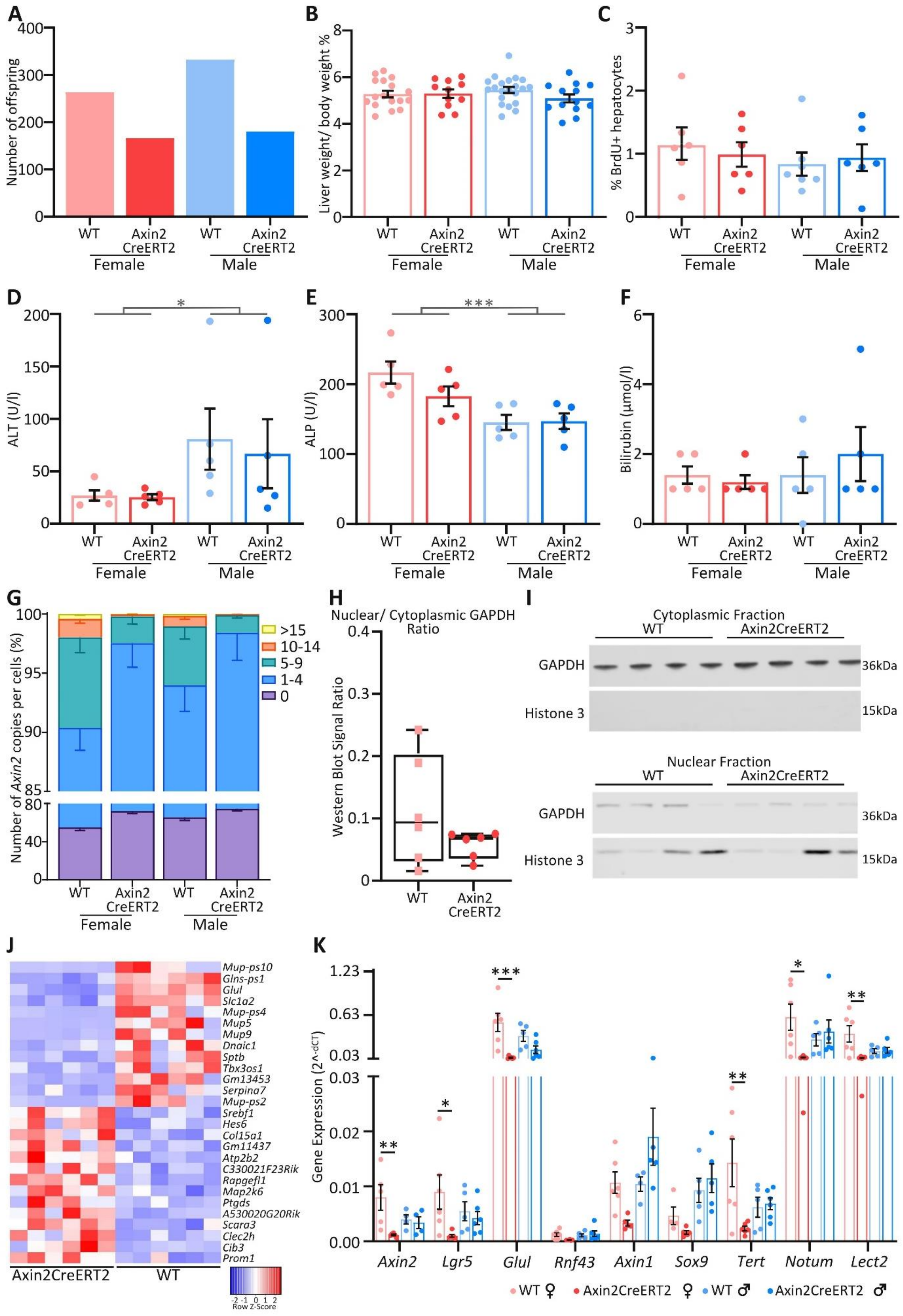

A
Number of offspring
400
300
200
100
0
WT
Axin2 CreERT2
Female
Male
B
Liver weight/ body weight %
8
6
4
2
0
C
% BrdU+ hepatocytes
3
2
1
0
D
ALT (U/l)
200
150
100
50
*
E
ALP (U/l)
300
***
F
Bilirubin (μmol/l)
G
Number of Axin2 copies per cells (%)
>15
10-14
5-9
1-4
0
H
Nuclear/ Cytoplasmic GAPDH Ratio
Western Blot Signal Ratio
0.4
0.3
0.2
0.1
I
Cytoplasmic Fraction
Nuclear Fraction
Axin2CreERT2
GAPDH
Histone 3
36kDa
15kDa
J
Mup-ps10
Glns-ps1
Glul
Slc1a2
Mup-ps4
Mup5
Mup9
Dnaic1
Sptb
Tbx3os1
Gm13453
Serpina7
Mup-ps2
Srebf1
Hes6
Col15a1
Gm11437
Atp2b2
C330021F23Rik
Rapgefl1
Map2k6
Ptgds
A530020G20Rik
Scara3
Clec2h
Cib3
Prom1
Axin2CreERT2
WT
-2 -1 0 1 2
Row Z-Score
K
Gene Expression (2^-dCT)
1.23
0.63
0.03
0.02
0.01
0.00
Axin2
Lgr5
Glul
Rnf43
Axin1
Sox9
Tert
Notum
Lect2
WT ♀
Axin2CreERT2 ♀
WT ♂
Axin2CreERT2 ♂

#### Sup Fig 3: Altered physiology in mice carrying the Axin2CreERT2 allele

(**A**) Number of WT and Axin2CreERT2$^{+/WT}$ mice born from WT x Axin2CreERT2$^{+/WT}$ breeding pairs split by sex (Female: WT = 263, Axin2CreERT2$^{+/WT}$ = 166, Male: WT = 332, Axin2CreERT2$^{+/WT}$ = 180). (**B**) Liver weight to body weight ratio of uninduced WT and Axin2CreERT2$^{+/WT}$ mice; N = 21/13 and 17/11 in WT /Axin2CreERT2$^{+/WT}$ mice for males and females respectively, two-way ANOVA, mean ± SEM. (**C**) Quantification of homeostatic hepatocyte proliferation in uninduced WT and Axin2CreERT2$^{+/WT}$ mice given BrdU 2hrs prior to harvest from IHC sections stained with BrdU; N = 6 mice per cohort except WT male where N = 7 per group, two-way ANOVA, mean ± SEM. Blood plasma analysis of (**D**) alanine transaminase (ALT), (**E**) alkaline phosphatase (ALP) and (**F**) total bilirubin from uninduced WT and Axin2CreERT2$^{+/WT}$ mice; N = 5 mice, two-way ANOVA, mean ± SEM. (**G**) Quantification of the number of *Axin2* mRNA copies per hepatocyte from sections probed using RNA *in situ* hybridisation for *Axin2*; N = 6 mice per cohort, mean ± SEM. (**H**) Quantification of the nuclear to cytoplasmic ratio of GAPDH in protein fractions from uninduced female WT and Axin2CreERT2$^{+/WT}$ livers. Protein fractions were deemed contaminated when the signal ratio was above 0.2; N = 6 mice per cohort, t-test, box plot with centre line showing the median values (interquartile range (IQR)), for WT 0.09 (0.03-0.20) and Axin2CreERT2$^{WT/-}$ 0.07 (0.04-0.07), whiskers represent minimum and maximum values. (**I**) Representative western blots displaying successful fractionation of liver tissue as shown by the depletion of Histone 3 in the cytoplasmic protein fraction and an enrichment of Histone 3 in the nuclear fraction from uninduced female WT and Axin2CreERT2$^{+/WT}$ mice. In the cytoplasmic fraction, GAPDH is used as a loading control, however in the nuclear fraction GAPDH is used as an integrity control. Note: the GAPDH western blot in Sup Fig. 3I is the same blot that is used in Fig. 2G, but with a shorter exposure. (**J**) Heat map displaying significant differentially expressed genes identified by RNAseq analysis of whole female livers in uninduced Axin2CreERT2$^{+/WT}$ compared to WT mice (N = 6). (**K**) Gene expression analysis of Wnt target genes from whole livers by qRT-PCR of uninduced WT and Axin2CreERT2$^{+/WT}$ mice split by sex; normalised female data shown in Fig. 2n; N = 4-6, two-way ANOVA, mean ± SEM. P = *<0.05; **<0.01; ***<0.001; **** <0.0001. Statistical comparisons between female vs. male are reported as the interaction factors from two-way ANOVA.

| Gene Name | Catalogue number | Probe Name |
|---|---|---|
| *Axin2* | QT00126539 | Mm_Axin2_1_SG QuantiTect Primer Assay |
| *Lgr5* | QT00123193 | Mm_Gpr49_1_SG QuantiTect Primer Assay |
| *Glul* | QT01062306 | Mm_Glul_1_SG QuantiTect Primer Assay |
| *Rnf43* | QT01777251 | Mm_Rnf43_1_SG QuantiTect Primer Assay |
| *Axin1* | QT00249494 | Mm_Axin1_1_SG QuantiTect Primer Assay |
| *Sox9* | QT00163765 | Mm_Sox9_1_SG QuantiTect Primer Assay |
| *Tert* | QT00104405 | Mm_Tert_1_SG QuantiTect Primer Assay |
| *Notum* | QT01749559 | Mm_Notum_2_SG QuantiTect Primer Assay |
| *Lect2* | QT00113554 | Mm_Lect2_1_SG QuantiTect Primer Assay |
| *E-cadherin* | QT00121163 | Mm_Cdh1_1_SG QunatiTect Primer Assay |
| *Rn18s* | QT02448075 | Mm_Rn18s_3_SG QuantiTect Primer Assay |

Sup Table 1: List of primers used for qRT-PCR

### Methods

#### Animal Models

All animal experiments were performed in accordance with a UK Home Office project licence (70/8891; protocol numbers 2, 3 and 4), in accordance with Home Office and ARRIVE guidelines[1] and were subject to review by the animal welfare and ethical review board of the University of Glasgow. To minimise pain, suffering and distress to the animals, single use needles and non-adverse handling techniques were used throughout. The animals used in this study were on a C57BL6/J background. Both male and female mice were housed in a specific pathogen-free environment and kept under standard conditions with a 12 hr day/night cycle with access to food and water *ad libitum*. Environmental enrichments, in the form of bedding, plastic tunnels and chew sticks, were added to all cages. The following transgenic mouse strains were used: Axin2CreERT2[2], R26R-LSL-tdTomato[3] (herein referred to as LSL-RFP) and Tcf/Lef:H2B-GFP[4]. Genotyping of ear notches taken at weaning (3 weeks of age) was performed by Transnetyx.

All mice used in this study were either induced or sampled (uninduced) between 8-12 weeks of age, unless otherwise specified. All mice were age and sex matched littermates and were administered BrdU (250µl, GE Healthcare) via intraperitoneal (i.p.) injection 2hrs prior to culling unless otherwise specified. BrdU in drinking water (0.8 mg/ml) was administered to female Axin2CreERT2$^{WT/WT}$; Tcf/Lef:H2B-GFP$^{+/-}$ and Axin2CreERT2$^{+/WT}$; Tcf/Lef:H2B-GFP$^{+/-}$ mice after 70% partial hepatectomy for 2 days. The animal studies were not randomised and the experimenters were not blinded to the experiments as induced mice with the LSL-RFP reporter develop red skin, however analysis was performed using standardised automated analyses where possible. All mice were health checked daily.

For wild-type analysis, C57BL6/J mice were either born in house (confirmed ≥ N10 by SNP analysis (Taconic.com)) or supplied by Charles River Laboratories and were allowed to acclimatise for at least 1 week prior to experimentation. C57BL6/J mice of both sexes (9 weeks old) received a single i.p. injection of 4mg tamoxifen (10 mg/ml stock, Sigma-Aldrich), dissolved in 10% ethanol/ 90% corn oil (Sigma) and were sampled 2 days post induction[5]. Tamoxifen controls were uninduced litter mates. Mice for this study had mean (IQR(interquartile range)) body weights of 24.1g (23.7-24.8) and 18.8g (18.8-19.1) for males and females respectively at induction (including uninduced controls). For homeostatic proliferation studies, male and female wild-type mice between 8-12 weeks old (young) or between 12-14 months old (aged) were administered a single BrdU i.p. injection 2hrs prior to harvesting.

To assess the effect of the Axin2CreERT2 knock-in construct, tissue was harvested from uninduced male and female mice harbouring Axin2CreERT2$^{WT/WT}$; Tcf/Lef:H2B-GFP$^{+/-}$ (control, herein referred to as WT) and Axin2CreERT2$^{+/WT}$; Tcf/Lef:H2B-GFP$^{+/-}$ alleles between 8-12 weeks of age.

For lineage tracing studies, Axin2CreERT2$^{+/WT}$; LSL-RFP$^{+/-}$ mice of both sexes (8-12 weeks of age) were induced with a single injection of 3mg or 4mg tamoxifen by i.p. injection. Mice were sampled at either 2, 5, 7, 200 or 365 days post induction. Mice for lineage tracing had mean (IQR) body weights of 26.7g (25.4-27.8) and 20.8g (19.3-21.7) for males and females respectively at induction.

To induce zonal liver injury, Axin2CreERT2$^{+/WT}$; LSL-RFP$^{+/-}$ female mice (11-22 weeks of age), were first induced by four consecutive daily i.p. injections of 3, 2, 2 and 2mg tamoxifen. On day 7 after the first tamoxifen injection, mice were either harvested (baseline) or received a single i.p. injection of carbon tetrachloride ($CCl_4$; Sigma), diluted 1:4 in corn oil (Sigma), at a dose of 1ul/g and were harvested at either 1, 2 or 4 days after $CCl_4$ administration.

70% Partial hepatectomy (PHx) was performed on 1) uninduced male and female WT and Axin2CreERT2$^{+/WT}$ mice and 2) uninduced female Axin2CreERT2$^{WT/WT}$; Tcf/Lef:H2B-GFP$^{+/-}$ and Axin2CreERT2$^{+/WT}$; Tcf/Lef:H2B-GFP$^{+/-}$ mice under sevoflurane anaesthesia (with $O_2$) in a sterile surgical field as previously described with minor modifications[6]. Briefly, a median laparotomy was performed followed by ligation and removal of the left lateral and median lobes; the right and left portions of the median lobe were ligated separately to preserve the biliary tree and gallbladder. Pre and post-operative analgesia were administered (0.1 mg/kg buprenorphine (pre) and 0.05 mg/kg buprenorphine (post)), clear $H_2O$ diet gel packs (Tecniplast) were also added to the cage during recovery. Mice were harvested either 2 or 7 days after surgery and were administered BrdU 2hrs prior to sampling, or for 2 days in the drinking water (0.8 mg/ml). Mice had mean (IQR) body weights of 25.9g (25.0-26.8) and 20.9g (19.7-22.1) for males and females respectively at surgery.

All mice were euthanized by $CO_2$ inhalation. Blood was collected by cardiac puncture, placed in lithium heparin collection tubes (Sarstedt) and centrifuged (2000g for 10 mins) and the plasma separated prior to storage at -80°C. The liver was harvested and fixed in 10% neutral buffered formalin (in PBS) for 24hrs whilst the caudate lobe was immediately frozen on dry ice and stored at -80℃. Formalin-fixed tissue was then embedded into paraffin blocks.

Power calculations were not routinely performed; however, animal numbers were chosen to reflect the expected magnitude of response taking into account the variability observed in pilot experiments using mice induced outside the 8-12 week age induction window (additional data from these mice available upon request). For all experiments the number of biological replicates ≥ 3 mice per cohort.

#### Liver Biochemistry

Plasma samples were analysed for alanine transaminase (ALT), alkaline phosphatase (ALP) and total bilirubin using the Dimension Expand chemistry system (Siemens Healthcare Diagnostics), following IFCC (International Federation of Clinical Chemistry) approved methods.

#### Immunohistochemistry (IHC), Immunofluorescence (IF), and *in situ* hybridization (ISH)

Standard immunohistochemistry techniques were utilised throughout this study, detailed protocols are available upon request. Staining was performed on 4µm sections. The following primary antibodies (dilutions) were used: RFP (1/1000 (IHC) or 1/200 (IF); 601-401-379, Tebu-bio), GS (1/600 (IHC); HPA007316, Sigma Aldrich or 1/1000 (IF); 610517, BD Biosciences), HNF4α (1/40 (IF); SC6556, Santa Cruz Biotechnology), E-cadherin (1/500 (IF); 610181, BD Biosciences), BrdU (1/250 (IHC); 347580, BD Biosciences or 1/500 (IF); ab6326, AbCam) and GFP (1/600 (IHC); 2555, Cell Signalling). IHC detection was performed with DAB (DAKO) followed by counterstaining with Haematoxylin. IF detection used species specific Alexa Fluor 488, 555 or 647 probes (A21208/A21202, A31572/A31570/S32355, and A21447 (Invitrogen, UK), respectively. Nuclei were detected using DAPI (0100-20, SouthernBiotech). Negative controls excluding each/all primary antibodies were routinely performed.

RNA *in situ-hybridisation* detection for *Axin2 and Lgr5* was performed using RNAscope 2.5 LS (Brown) detection kit (Advanced Cell Diagnostics, Hayward, CA; 322100; probes 400338 and 312178 respectively) on a Leica Bond Rx autostainer strictly according to the manufacturer's instructions. The quality and integrity of RNA in the tissues was evaluated using a positive control probe (*Ppib*; 313918).

#### Image analysis

IHC and ISH images were digitalised using a Leica Aperio AT2 slide scanner (Leica Microsystems, UK) at 20x magnification. Scanned images were analysed blindly using HALO Image analysis software (V3.1.1076.363, Indica Labs). Liver boundaries were manually defined using the HALO software. Classifiers were applied to identify hepatocytes, and livers were analysed for the percentage of positive hepatocytes for the nuclear stains GFP and BrdU, whereas percentage of positive tissue area were analysed for the cytoplasmic stains RFP and GS. For quantification of ISH, the number of probe copies and hepatocytes were determined.

Fluorescent tiled images were generated on an Opera Phenix High-Content Screening System (Perkin Elmer) at 20x magnification. DAPI, Alexa Fluor 488, 555 and 647 were detected using excitation wavelengths of 405, 488, 561 and 640nm, and emission band paths of 435-480, 500-550, 570-630 and 650-760nm, respectively using the same settings throughout. Consecutive non-overlapping fields were analysed blindly using Columbus Image analysis software (2.8.0.138890, Perkin Elmer). Positivity gating thresholds were defined using negative controls. For representative images, processing adjustments were performed equally.

##### Quantitative reverse transcription polymerase chain reaction (qRT-PCR)

Total RNA was extracted from snap frozen liver tissue samples using the Qiagen RNeasy Mini kit according to the manufacturer's instructions, including the optional DNA degradation step (74104, Qiagen, UK). cDNA synthesis with the integrated removal of genomic DNA was performed using the QuantiTect® Reverse Transcription Kit (205313, Qiagen, UK) on a PTC-200 thermal cycler (MJ Research), according to the manufacturer's instructions. qRT-PCR was performed using QuantiTect SYBR Green PCR Kit (204145, Qiagen, UK) and QuantiTect Primer Assays (249900, Qiagen, UK) following the two-step RT-PCR standard protocol (10µl final volume per well) on the QuantStudio™ 5 Real-Time PCR 384-well system (Applied Biosystems). Data were analysed following normalisation to the housekeeping gene 18S ribosomal RNA (*Rn18s* (Mouse)). All samples were run in triplicate, primer catalogue numbers are shown in Supplementary Table 1.

##### RNAseq Analysis

RNA from sex matched whole liver samples was extracted as described above. For partial hepatectomy the changes were compared between paired baseline (0hrs, resected tissue) and 48hrs after surgery from the same mouse. The quality of the purified RNA was tested on an Agilent 2200 Tapestation (D1000 screentape) using RNA screentape, RIN values of >7 were obtained throughout. Libraries for cluster generation and DNA sequencing were prepared following an adapted method[7] using the Illumina TruSeq Stranded mRNA LT Kit. The quality and quantity of the DNA libraries was assessed on an Agilent 2200 Tapestation and Qubit (Thermo Fisher Scientific) respectively. The libraries were run on the Illumina Next Seq 500 using the High Output 75 cycles kit (2x36cycles, paired end reads, single index).

Fastq files were generated from the sequencer output using illumina's bcl2fastq version 2.20.0.422 and quality checks on the raw data were done using FastQC version 0.11.7[8] and Fastq Screen version 0.14.0[9]. Alignment of the RNA-Seq paired-end reads was to the GRCh38[10] version of the mouse genome and annotation using Hisat2 version 2.1.0[11]. Expression levels were determined and statistically analysed by a workflow combining HTSeq version 0.11.2[12], the R environment version 3.6.1[13], utilising packages from the Bioconductor data analysis suite[14] and differential gene expression analysis based on the negative binomial distribution using the DESeq2 package[15]. Further data analysis and visualisation used R and Bioconductor packages. Pathway analysis and GO Processes were completed using the MetaCore software from Clarivate Analytics. Gene Set Enrichment Analysis (GSEA)[16] was used to determine whether a set of Wnt associated genes[17] were significantly different between the uninduced WT and Axin2CreERT2$^{+/WT}$ mice. Genes with at least 2 reads per million in at least one sample from any experimental group were included in the ranked comparisons. The data have been deposited in Geo at NCBI accession number GSE (accession number available upon acceptance).

##### Isolation of enriched cytoplasmic and nuclear fraction from liver tissue

Between 100–300mg of liver tissue from uninduced Axin2CreERT2$^{WT/WT}$; Tcf/Lef:H2B-GFP$^{+/-}$ and Axin2CreERT2$^{+/WT}$; Tcf/Lef:H2B-GFP$^{+/-}$ female mice was weighed and processed by pestle and mortar on liquid nitrogen. The resulting tissue powder was then resuspended in hypotonic buffer (10 mM HEPES pH 7.9, 10 mM KCl, 0.1 mM EDTA, 0.1 mM EGTA, 1x (w/v) protease inhibitor cocktail, 10 mM NaF, 1mM Na3VO4, 1 mM DTT, 1 mM PMSF). Following 15 mins incubation on ice, the suspension was vortexed at full speed for 10 secs and centrifuged (2300g, 5 mins). The supernatant was retained to represent the soluble cytoplasmic fraction. The resulting pellet was washed twice more with fresh hypotonic buffer and further supernatant discarded, to remove contaminating cytoplasmic fraction The subsequent enriched nuclear pellet was resuspended in high salt buffer (20 mM HEPES pH 7.9, 420 mM NaCl, 1.5 mM $MgCl_2$, 20% (v/v) glycerol, 1x (w/v) protease inhibitor cocktail, 10 mM NaF, 1 mM $Na_3VO_4$, 1 mM DTT, 1 mM PMSF) and incubated on ice for 15 mins with intermittent low speed vortexing to ensure homogeneity. Centrifugation was performed (15,000g, 30 mins, 4°C) to separate the soluble and insoluble enriched nuclear fractions. The supernatant containing enriched soluble nuclear fraction was collected and diluted with 2 volumes of dilution buffer (20 mM HEPES pH 7.9, 20% (v/v) glycerol, 0.5% (v/v) Igepal CA630). Laemlli buffer (x4) was added to both enriched cytoplasmic and enriched nuclear samples before boiling at 95°C.

##### Western Blot

Nuclear and cytoplasmic protein samples from WT and Axin2CreERT2$^{+/WT}$ mice were quantified using a Bradford assay (5000006, BioRad), according to the manufacturer's instructions. Equal amounts of protein were separated by Bis-Tris SDS-Page using NuPAGE pre cast 4-12% gels (NP0321BOX, Invitrogen) and transferred onto 0.25 µm nitrocellulose membrane. Total protein was visualised with Ponceau S. Membranes were blocked in 5% Milk-TBST and probed with antibodies. The following primary antibodies (1:500 dilution) were used: β-catenin (610153, BD Transduction Laboratories), histone 3 (ab1791, Abcam), β-actin (691001, MP Biomedicals), and GAPDH (SC-32233, Santa Cruz Biotechnology). Li-COR Bioscience IR-Dye-labelled secondary antibodies (680, 926-68070/926-68071 and 800, 926-32210/926-32211) were used for visualisation (1/10,000) using the Li-COR Odyssey Infrared Imager. Determination of β-catenin ratio between the enriched cytoplasmic and nuclear fraction was conducted by initial normalisation to fraction specific loading controls, GAPDH for enriched cytoplasmic fraction and β-actin for enriched nuclear fraction. A ratio between the normalised nuclear/ cytoplasmic β-catenin bands was calculated. Data were excluded from analysis when nuclear/ cytoplasmic GAPDH signal ratios were more than 0.2, inferring that the nuclear fraction was contaminated with cytoplasmic protein. Detailed protocols are available upon request.

##### Statistical analysis

Vehicle or genotype controls were used and no bias was applied during husbandry, during tissue sampling or during outcome analysis. GraphPad Prism software (version 8.4.3) was used for all statistical analyses. To determine whether data were normally distributed a Shapiro-Wilk normality test was used. Data with a normal distribution were analysed using a two-tailed unpaired t-test. For data with more than two groups a one-way ANOVA was used. For data where two groups are split between two independent variables, a two-way ANOVA was used. For data that did not follow a normal distribution because the N was too small, a two-tailed Mann–Whitney test was used unless otherwise stated. For correlation studies a linear regression was used. Data are presented as mean ± SEM.; N refers to the number of biological replicates and unless indicated otherwise, P values are ∗P < 0.05; ∗∗P < 0.01; ∗∗∗P <0 .001 and ∗∗∗∗P <0 .0001. Figures were constructed using Scribus v1.4.7 (GNU general public license), Photoshop v12.1 and Aperio ImageScope v12.4.0.5043.

#### Method References


1 Percie du Sert, N. *et al.* The ARRIVE guidelines 2.0: Updated guidelines for reporting animal research. *PLOS Biology* **18**, e3000410, doi:10.1371/journal.pbio.3000410 (2020).

2 van Amerongen, R., Bowman, A. N. & Nusse, R. Developmental stage and time dictate the fate of Wnt/β-catenin-responsive stem cells in the mammary gland. *Cell Stem Cell* **11**, 387-400, doi:10.1016/j.stem.2012.05.023 (2012).

3 Madisen, L. *et al.* A robust and high-throughput Cre reporting and characterization system for the whole mouse brain. *Nature Neuroscience* **13**, 133-140, doi:10.1038/nn.2467 (2010).

4 Ferrer-Vaquer, A. *et al.* A sensitive and bright single-cell resolution live imaging reporter of Wnt/ß-catenin signaling in the mouse. *BMC developmental biology* **10**, 121, doi:10.1186/1471-213x-10-121 (2010).

5 Wang, B., Zhao, L., Fish, M., Logan, C. Y. & Nusse, R. Self-renewing diploid Axin2(+) cells fuel homeostatic renewal of the liver. *Nature* **524**, 180-185, doi:10.1038/nature14863 (2015).

6 Mitchell, C. & Willenbring, H. A reproducible and well-tolerated method for 2/3 partial hepatectomy in mice. *Nature protocols* **3**, 1167-1170, doi:10.1038/nprot.2008.80 (2008).

7 Fisher, S. *et al.* A scalable, fully automated process for construction of sequence-ready human exome targeted capture libraries. *Genome Biology* **12**, R1, doi:10.1186/gb-2011-12-1-r1 (2011).

8 Andrews, S. *FastQC: a quality control tool for high throughput sequence data.*, <http://www.bioinformatics.bbsrc.ac.uk/projects/fastqc> (2018).

9 Wingett, S. *FastQ Screen*, <http://www.bioinformatics.babraham.ac.uk/projects/fastq_screen/> (2018).

10 Zerbino, D. R. *et al.* Ensembl 2018. *Nucleic Acids Res* **46**, D754-d761, doi:10.1093/nar/gkx1098 (2018).

11 Kim, D., Langmead, B. & Salzberg, S. L. HISAT: a fast spliced aligner with low memory requirements. *Nature Methods* **12**, 357-360, doi:10.1038/nmeth.3317 (2015).

12 Anders, S., Pyl, P. T. & Huber, W. HTSeq--a Python framework to work with high-throughput sequencing data. *Bioinformatics (Oxford, England)* **31**, 166-169, doi:10.1093/bioinformatics/btu638 (2015).

13 R Team*: A language and environment for statistical computing. R Foundation for Statistical Computing, Vienna, Austria.* , <https://www.R-project.org/> (2018).

14 Huber, W. *et al.* Orchestrating high-throughput genomic analysis with Bioconductor. *Nature methods* **12**, 115-121, doi:10.1038/nmeth.3252 (2015).

15 Love, M. I., Huber, W. & Anders, S. Moderated estimation of fold change and dispersion for RNA-seq data with DESeq2. *Genome Biology* **15**, 550, doi:10.1186/s13059-014-0550-8 (2014).

16 Subramanian, A. *et al.* Gene set enrichment analysis: A knowledge-based approach for interpreting genome-wide expression profiles. *Proceedings of the National Academy of Sciences* **102**, 15545, doi:10.1073/pnas.0506580102 (2005).

17 Sansom, O. J. *et al.* Loss of Apc in vivo immediately perturbs Wnt signaling, differentiation, and migration. *Genes and Development* **18**, 1385-1390 (2004).
